# A Chromosome-level Genome Assembly of the Potato Leafhopper *Empoasca fabae* (Hemiptera: Cicadellidae)

**DOI:** 10.64898/2026.07.04.736200

**Authors:** Joshua Molligan, Florent Sylvestre, Edel Pérez-López

## Abstract

The potato leafhopper, *Empoasca fabae* (Harris, 1841), is a highly polyphagous, migratory insect pest of eastern North America that feeds on more than 200 herbaceous and woody plant species, causing substantial losses to forage and field crops. Despite its agricultural and ecological importance, no genome has been available for this species. Here, we present the first chromosome-level genome assembly of *E. fabae*, generated from Oxford Nanopore long reads, Illumina short reads, and Omni-C proximity-ligation data. The final assembly spans 908 Mb across 132 scaffolds, with 99.8% of the assembly captured in ten chromosome-length scaffolds (nine autosomes and an X chromosome) with a scaffold N50 of 96.2 Mb. The assembly is highly complete, recovering 92.4% of conserved hemipteran single-copy orthologs, and is composed of 47.6% repetitive sequence, dominated by long terminal repeat retrotransposons and unclassified elements. Read-depth comparison between male and female individuals supports assignment of a single sex-linked chromosome, consistent with an XO sex-determination system. BRAKER3 gene annotation predicted 31,406 protein-coding genes after retaining the longest isoform per locus. Comparative genome analysis against the two closest related Typhlocybinae species with genomes available, *Matsumurasca onukii* and *Hebata decipiens*, revealed extensive chromosome-scale collinearity, while defining a shared core gene repertoire. This reference genome provides a foundation for comparative and population genomic studies and for investigating genetic traits in this economically important crop pest species.

**SIGNIFICANCE:** Leafhoppers (Cicadellidae) are among the most diverse families of plant-feeding insects, but chromosome-level genomes remain scarce, particularly for mesophyll-feeding members of the subfamily Typhlocybinae. The potato leafhopper, *Empoasca fabae*, is an unusually polyphagous crop and migratory pest of major importance across North America. Here, we provide the first chromosome-level genome assembly for this species. This chromosomal reference reveals broad synteny with two related Typhlocybinae relatives. This assembly will serve as a critical resource, enabling further comparative genomics, population genomics, and functional studies of host-plant adaptation in a significant agricultural crop pest species.

## INTRODUCTION

The potato leafhopper, *Empoasca fabae* (Harris, 1841), is a migratory, polyphagous insect pest native to eastern North America. It is reported to feed on over 200 herbaceous and woody plant species and is known to cause injury to numerous economical crops, such as alfalfa, potato and soybean (Lamp et al., 1994). Unlike most leafhoppers, which feed from phloem or xylem, *E. fabae* belongs to the mesophyll-feeding subfamily Typhlocybinae, whose members are known for rupturing parenchymal cells during feeding (Backus, 1988; Backus & Hunter, 1989). This cell-rupturing habit produces characteristic ‘hopperburn’ symptoms that drives much of the crop loss attributed to this species. *E. fabae* compounds this damage with the ability to vector plant pathogens and annual long-distance migrations. *E. fabae* is known to primarily overwinter in the southern United States with annual northern migrations each spring, reaching as far as Québec, Canada (Baker et al., 2015; Taylor & Shields, 2018; Santos et al., 2025). Together, its broad host range, feeding habits, and seasonal migration make *E. fabae* a compelling system for studying pest adaptation to new agricultural systems and climatic extremes.

Leafhoppers are among the most diverse families of plant-feeding insects, with more than 20,000 described species (Bartlett et al., 2018), yet chromosome-level genomic resources remain scarce relative to that diversity. The gap is especially striking for Typhlocybinae, given the agricultural importance of several of its members. Genomes from related species, including *Matsumurasca onukii* (Matsuda, 1952), a major pest of tea in East Asia, and *Hebata decipiens* (Paoli, 1930), offer useful comparative references to *E. fabae* (Raupach et al., 2002; Zhang et al., 2019), but no genome assembly has been available for *E. fabae* to date. Such a resource is needed to resolve genome organization, repetitive DNA content, and sex-chromosome structure, in addition to aiding future comparative and population genomic studies of this economically important pest.

Here, we present the first chromosome-level and species reference genome assembly of *E. fabae*, generated from Oxford Nanopore long reads, Illumina short reads, and Omni-C proximity-ligation data. We resolve ten chromosome-length scaffolds, nine autosomes and one X chromosome, and annotate the repeat landscape and protein-coding gene repertoire. We then compare genome structure and gene content against two related Old World Typhlocybinae. This assembly provides a foundational resource for future studies of host-plant adaptation, migration, and pest biology in *E. fabae* and related leafhoppers.

## RESULTS AND DISCUSSION

### A Chromosome-level Assembly of the *Empoasca fabae* Genome

High-molecular-weight genomic DNA was extracted from field-collected male *Empoasca fabae* (**Figure 1A**) and sequenced using Oxford Nanopore long-read and Illumina short-read technologies, complemented by an Omni-C proximity-ligation library for chromosome-scale scaffolding. The Nanopore long-read dataset (143 Gb) was read-corrected and assembled into a draft, from which alternative haplotypes were then removed, and subsequently polished twice with Illumina paired-end reads (160 Gb). Contamination screening of the draft assembly with FCS-GX removed any contigs assigned to non-target taxa. Chromosomal scaffolding of the draft assembly using Omni-C reads (300 Gb), followed by manual curation, produced the final assembly with a clear and well-resolved contact map (**Figure 1B**).

**Figure 1.**
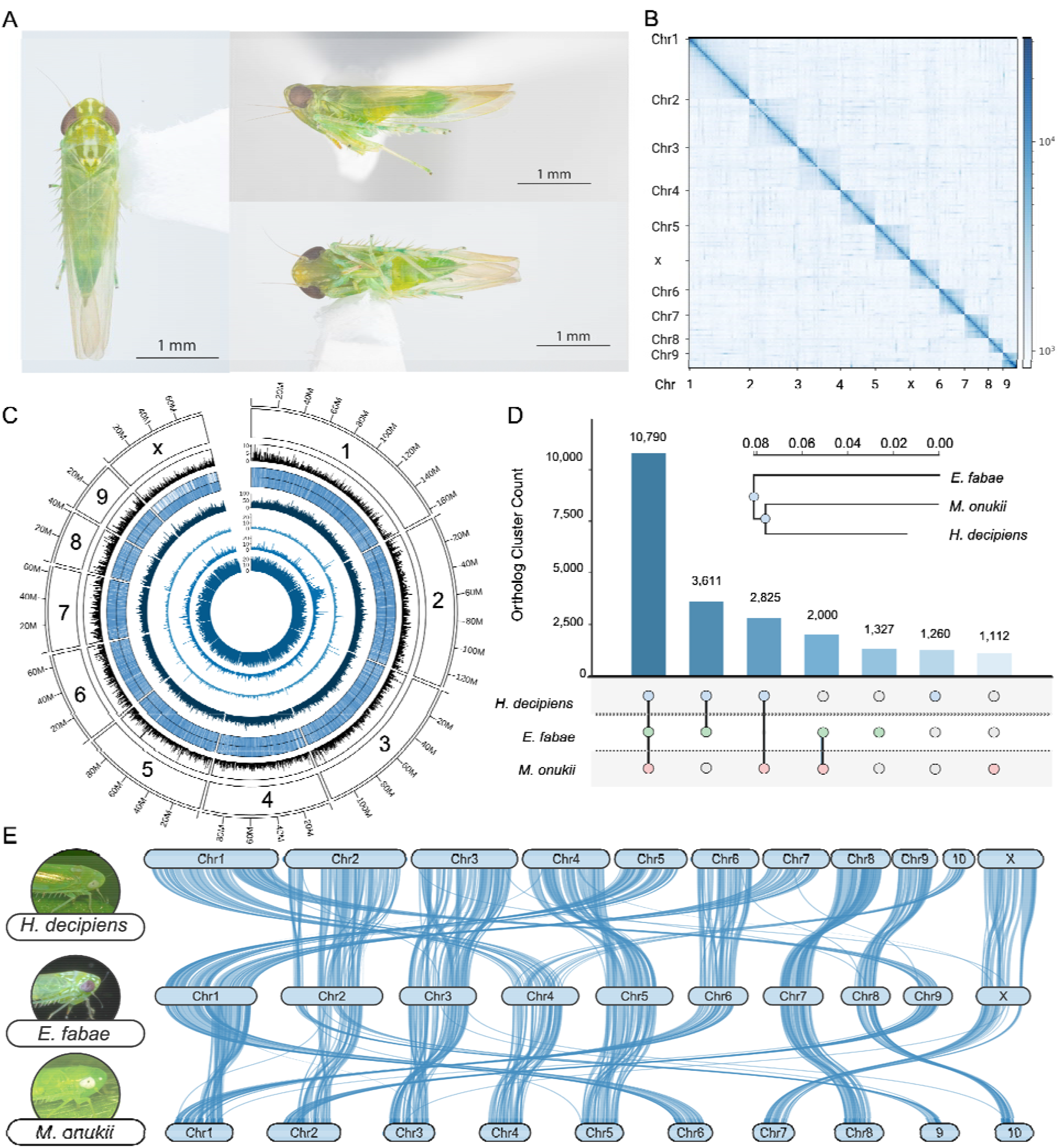
Chromosome-level genome assembly and comparative genomics of *Empoasca fabae*. (**A**) Specimen photographs of *E. fabae* males in dorsal, lateral, and ventral poses. Photographs credited to Jordanne Jacques, from EdeLab. (**B**) Omni-C contact map of the final scaffolded assembly, showing ten chromosome-length scaffolds (nine autosomes and the X chromosome). (**C**) Circos diagram of the ten chromosomes, with tracks from outer to inner representing: chromosome ideogram with scale in Mb; gene density in 100 kb windows, scaled 0-10; read coverage of male (outer) and female (inner) short-read libraries in 10 kb sliding windows; and repeat density in 100 kb windows for LINE, LTR, DNA, and simple repeats. (**D**) UpSet plot of orthogroup sharing among *E. fabae, Matsumurasca onukii*, and *Hebata decipiens*. Species tree is based on orthologous protein clusters. (**E**) Chromosome-scale synteny among the three genomes, with collinear blocks of single-copy orthologs shown as connecting ribbons. Photographs of *H. decipiens* (top), *E. fabae* (middle), and *M. onukii* (bottom) obtained from Wikimedia Commons and credited to Hectonichus (CC BY-SA 3.0), Xpda (CC BY-SA 4.0), and Christian Ferrer (CC BY 4.0), respectively.

The final assembly spans 908 Mb across 132 scaffolds, with a GC content of 35.9% (**Table 1**). The majority of the assembly is in ten chromosome-length scaffolds, comprising nine autosomes and an X chromosome (**Figure 1C**). The largest of these reaches 164 Mb, and the smaller, residual scaffolds together account for only a fraction of total length. Mean read depth across chromosomes was 32.7x, supporting a uniform coverage. At the scaffold level the assembly is highly contiguous, with a scaffold N50 of 96.2 Mb, an N90 of 40.9 Mb, and L50 and L90 values of 4 and 9, respectively (**Table 1**).

**Table 1.** Genome assembly statistics for the *Empoasca fabae* reference genome.

| Metric | Scaffolds | Contigs |
| --- | --- | --- |
| Number of contigs (≥ 1,000 bp) | 132 | 10,582 |
| Number of contigs (≥ 5,000 bp) | 132 | 10,531 |
| Number of contigs (≥ 10,000 bp) | 132 | 10,476 |
| Number of contigs (≥ 25,000 bp) | 105 | 8,166 |
| Number of contigs (≥ 50,000 bp) | 67 | 5,293 |
| Total length (≥ 1,000 bp) | 908,556,302 bp | 903,314,055 bp |
| Total length (≥ 5,000 bp) | 908,556,302 bp | 903,158,603 bp |
| Total length (≥ 10,000 bp) | 908,556,302 bp | 902,753,937 bp |
| Total length (≥ 25,000 bp) | 908,032,845 bp | 857,615,848 bp |
| Total length (≥ 50,000 bp) | 906,664,310 bp | 754,825,845 bp |
| Largest contig | 163,687,506 bp | 1,413,263 bp |
| Total assembly length | 908,556,302 bp | 903,321,273 bp |
| GC content (%) | 35.91 | 35.91 |
| N50 | 96,186,748 bp | 141,938 bp |
| N90 | 40,996,017 bp | 35,464 bp |
| L50 | 4 | 1,728 |
| L90 | 9 | 6,669 |
| N's per 100 kbp | 576.13 | 0.00 |

Prior to scaffolding, the draft ONT assembly comprised 10,582 contigs with a contig N50 of 141.9 kb and a largest contig of 1.41 Mb (**Table 1**). The increase in contiguity from the contig to the scaffold level, together with a low gap content of approximately 575 N’s per 100 kb introduced during Omni-C scaffolding, indicates that the long-range proximity-ligation information used here was effective in orienting contigs into chromosome-length scaffolds. Assessment with benchmarking universal single-copy orthologs (BUSCO) against the hemiptera_odb10 lineage (2,510 orthologs) recovered 92.4% as complete, partitioned into 90.4% single-copy and 2.0% duplicated, with 2.7% fragmented and 4.9% missing (**Supplementary Table S1**). The low proportion of duplicated orthologs is consistent with effective removal of alternative haplotypes during assembly, with a small fraction of complete orthologs flagged due to internal stop codons (3.3%).

Repeat annotation showed that 47.6% of the genome (433 Mb) is composed of repetitive elements, with interspersed repeats accounting for the large majority (46.3%) (**Figure 1C, Supplementary Table S2**). Retroelements occupy 15.4% of the genome, dominated by LTR elements (9.1%), principally Gypsy/DIRS1 (4.5%) with smaller contributions from BEL/Pao (1.2%) and Ty1/Copia (0.6%), followed by LINEs (6.3%). DNA transposons represent a comparatively minor fraction (1.1%) and rolling-circle Helitrons are rare (0.02%). The single largest component, however, is a substantial number of unclassified repeats (29.8%), exceeding all other annotated TE classes combined. This abundance of unclassifiable repetitive sequence is characteristic of large hemipteran genomes which lack closely related, curated repeat libraries, and accounts for a significant portion of the overall genome size.

To identify the sex chromosome, we further sequenced both male and female individuals and mapped short-read datasets back to the assembly to compare normalized read depth across chromosomes (**Figure 1C, Supplementary Table S3**). We suspected an XO sex-determination system, as similarly reported for other leafhoppers, which can be observed in a shift in relative female coverage for an X chromosome present in two copies, as there is only a single copy in males. Nine autosomes showed similar female-to-male coverage ratios ranging from 0.83 to 0.86 (mean 0.85), whereas a single chromosome exhibited elevated female coverage with a raw female-male ratio of 1.09 (normalized to a ratio of 1.28, against an autosomal baseline of 1). As this ratio is not exactly double the autosomal baseline, we then further assessed synteny among *E. fabae* and closely related species to provide further support of sex chromosome assignment.

### Comparative Genomics and Chromosome-scale Synteny

To place the *E. fabae* genome in a comparative context, we annotated the publicly available genomes of two related Typhlocybinae species, *Matsumurasca onukii* (GCA_018831715.1) and *Hebata decipiens* (GCA_964267455.1) in addition to *E. fabae*, using identical workflows and excluding transcriptomic evidence to minimize transcriptome-driven annotation bias. Following repeat masking, protein-coding genes were annotated with BRAKER3 using Arthropoda protein evidence, retaining only the longest isoform of each gene. Resulting annotations contained 31,406, 32,525 and 65,729 longest isoforms for *E. fabae, M. onukii* and *H. decipiens*, respectively. Orthogroup clustering across the three genomes identified 10,790 orthogroups shared by all three species, defining the core gene repertoire of this group (**Figure 1D**). As the observed variation in species orthogroup numbers could have been influenced by annotation quality or assembly fragmentation, we then emphasize using only shared core orthogroups and single copy orthologs for synteny and phytogenic inference rather than interpreting unique orthogroups for definitive gene gains or losses (**Figure 1D**). The resulting orthogroup-based species tree recovered *M. onukii* and *H. decipiens* as more closely related to each other than to *E. fabae*, consistent with the geographic and ecological separation of these taxa.

To examine chromosome-scale conservation, we extracted single-copy orthologs (7,169 per species) and identified collinear blocks among the three genomes using MCScanX (**Figure 1E**). This analysis assigned 17,971 of 21,507 genes (83.6%) to collinear blocks, distributed across 991 blocks and comprising 14,756 anchored gene pairs. Collinear blocks were substantial, with a median of 11 anchored genes per block and the largest spanning 174 consecutive anchors. The degree of synteny observed mirrored phylogenetic placement. The two Old World Typhlocybinae species, *M. onukii* and *H. decipiens*, shared the most extensive collinearity (6,204 anchored gene pairs, 87% of single-copy genes; median block size 14, largest block 174 anchors), whereas the comparisons against *E. fabae* recovered fewer and smaller blocks (4,361 and 4,191 anchored pairs, 58% and 61% of single-copy genes; median block sizes 9 and 10, compared to either *M. onukii* or *H. decipiens*, respectively).

Collinearity provided support for the assignment of the X chromosome. The chromosome assigned as the X in *E. fabae* based on female-male coverage ratios was highly syntenic with a single chromosome in each of the other two genomes (**Figure 1D**). It anchored to *H. decipiens* ChrX through 224 gene pairs across 20 collinear blocks, and to a single *M. onukii* chromosome (Chr7) through 202 gene pairs across 19 blocks. The same conserved linkage group was recovered independently, where H. *decipiens* ChrX mapped to *M. onukii* Chr7 through 288 gene pairs. Together, both female-male coverage bias and conserved synteny is supportive of sex-chromosome assignment.

These results establish a contiguous chromosome-level reference genome for *E. fabae*, resolving sex chromosome assignment and demonstrating collinearity with two closely related leafhoppers. As the first chromosome-level assembly for this species, this genome fills an important gap in the genomic sampling of Cicadellidae and provides a foundation for future comparative, population, and functional genomic investigations for this distinctive insect pest.

## MATERIALS AND METHODS

### Insect Collection and DNA Extraction

Adult *E. fabae* were collected alive in July 2024 from the Capitale-Nationale region of southern Québec, Canada, using an entomological net. Morphological identification of male genitalia was performed as described in previous studies (Molligan et al., 2025; Plante et al., 2024). Specimens from field collection are deposited at the Canadian National Collection of Insects, Arachnids, and Nematodes, under the voucher numbers CNC2098398-2098407, with Dr. Joel Kits as the person responsible of the Hemiptera division. High-molecular-weight genomic DNA was extracted from these field-captured male *E. fabae* using a Nanobind CBB kit (PacBio, California, USA) according to the manufacturer’s instructions for insects for both long- and short-read sequencing data sets. From the same collection event, 200 mg of male adults were also used to generate an Omni-C proximity-ligation library (Dovetail Genomics, California, USA).

### Whole-Genome Sequencing

Genomic DNA was sequenced using Oxford Nanopore long-read technology (ONT) and Illumina paired-end sequencing (Plasmidsaurus, California, USA), yielding 143 Gb of ONT long reads and 160 Gb of Illumina data. The Omni-C library was sequenced to generate 300 Gb of proximity-ligation reads for chromosomal scaffolding. Two additional Illumina short-read datasets (both 9 Gb) were generated using the same protocols from both male and female individuals for sex-chromosome identification.

### Genome Assembly and Scaffolding

The ONT long-read dataset was first read-corrected using Canu v2.3 (Koren et al., 2017) and then assembled with WTDBG2 v2.5 (Ruan & Li, 2020). Alternative haplotypes were removed using Purge_haplotigs v1.1.3 (Roach et al., 2018). Illumina paired-end reads were used to polish the draft assembly with Pilon v1.24 (Walker et al., 2014). Contigs identified as contamination were removed from the draft assembly using FCS-GX (Astashyn et al., 2024). Chromosomal scaffolding was carried out using Omni-C reads and the 3D-DNA pipeline (Dudchenko et al., 2017), followed by manual curation and visualization in Juicebox (Dudchenko et al., 2018; Durand et al., 2016). Assembly statistics and completion were computed with QUAST v5.3.0 and BUSCO v5.8.2 (Gurevich et al., 2013; Simão et al., 2015).

### Repeat Annotation

Repetitive elements were identified using a *de novo repeat libr*ary generated with RepeatModeler v2.0.7 (Hubley et al., 2023) and masked using RepeatMasker v4.1.8 (Smit et al., 2013). Repeat categories are reported summarized by class and by proportion of the genome occupied.

### Sex-Chromosome Identification

The two short-read Illumina datasets generated from male and female insects were mapped to the scaffolded assembly using bwa-mem2 v2.3 (Vasimuddin et al., 2019), and per-base coverage was quantified with SAMtools v1.17 (Li et al., 2009). Normalized female and male depth profiles were compared across chromosomes to distinguish autosomes from the X chromosome, with a chromosome showing elevated relative female coverage assigned as the X, consistent with an expected XO sex-determination system.

### Gene Annotation, Comparative Genomics and Synteny Analysis

To ensure standardized and directly comparable gene models across species, genomes of *E. fabae* (GCA_057929975.1), *M. onukii* (GCA_018831715.1), and *H. decipiens* (GCA_964267455.1) were annotated under an identical workflow. For each species, the genome assembly was independently repeat-masked using a species-specific library generated with RepeatModeler v2.0.7, followed by masking with RepeatMasker v4.1.8. The masked assemblies were then provided as input to BRAKER3 v3.0.8 (Gabriel et al., 2024). For comparative annotations, BRAKER3 was run using a uniform evidence set that excluded RNA-seq data to avoid expression-driven biases among species. Instead, gene prediction relied exclusively on the shared Arthropoda reference proteome, supplied to GeneMark-ETP v.1.0 (Bruna et al., 2024) and GeneMark-EP+ v4.7.1 (Bruna et al., 2020) for protein-guided training, combined with AUGUSTUS v3.5.0 for final model generation. Only the longest isoform per locus was retained for downstream analyses.

Orthogroups were inferred with OrthoFinder v3.1.0 (Emms et al., 2025), which provided a classification of shared and species-specific genes across the three genomes. The resulting orthogroup protein sets were then imported into OrthoVenn3 v3.1.0 (Sun et al., 2023) for visualization. For synteny and chromosome-scale collinearity, orthologous genes were extracted and used as inputs for MCScanX v1.0 (Wang et al., 2012). MCScanX v1.0 was then used to identify collinear blocks and anchor gene pairs across the three genomes. Chromosome-level collinearity visualizations were made with SynViso (Bandi & Gutwin, 2020).

## ACKNOWLEDGEMENTS

We are thankful to resources provided by Université Laval’s high-performance computing infrastructure at the Institute of Biology Intégrative et des Systèmes (IBIS). We are also thankful for the displayed specimen photographs taken by Jordanne Jacques, to Abrãao Almeada Santos who helped collect insects for sequencing, and to Florent Sylvestre for valuable discussion and feedback concerning assembly methods. Figures were assembled in BioRender.com.

## AUTHOR CONTRIBUTIONS

Conceptualization, J.M. and E.P.-L.; Methodology, J.M.; Investigation, J.M. and E.P.-L.; Data curation and formal analysis, J.M.; Visualization, J.M.; Resources and funding acquisition, E.P.-L.; Project administration and supervision, E.P.-L.; Writing original draft, J.M. and E.P.-L; All authors contributed to review & editing of this manuscript.

## DATA AVAILABILITY

The *Empoasca fabae* genome has been deposited in the NCBI database under the accession number GCA_057929975.1 (BioProject ID PRJNA1435210), and is currently the reference genome for the species. The NCBI sequencing datasets generated for this study are publicly available through NCBI under the following SRA accessions: SRX32526771 (Short reads for *Empoasca fabae*), SRX32526772 (Proximity ligation data for *Empoasca fabae*), SRX32526773 (Long read data for *Empoasca fabae*), SRX32839852 (*Empoasca fabae* female) and SRX32839853 (*Empoasca fabae* male). *De novo annotation* files use for comparative genomics and chromosome-scale synteny of all three genomes are available on Zenodo: https://doi.org/10.5281/zenodo.21110129. The genome analysis workflow is available on GitHub: https://github.com/Edelab/Genome_assembly_empoasca_fabae.

## LITERATURE CITED

Astashyn A, Tvedte ES, Sweeney D, Sapojnikov V, Bouk N, Joukov V, Mozes E, Strope PK, Sylla PM, Wagner L, et al. Rapid and sensitive detection of genome contamination at scale with FCS-GX. Genome Biol. 2024:25(1):60. 10.1186/s13059-024-03198-7.

Backus EA. Sensory systems and behaviours which mediate hemipteran plant-feeding: a taxonomic overview. J Insect Physiol. 1988:34(3):151–165. 10.1016/0022-1910(88)90045-5.

Backus EA, Hunter WB. Comparison of feeding behavior of the potato leafhopper *Empoasca fabae (Homoptera*: Cicadellidae) on alfalfa and broad bean leaves. Environ Entomol. 1989:18(3):473–480. 10.1093/ee/18.3.473.

Baker MB, Venugopal PD, Lamp WO. Climate change and phenology: *Empoasca fabae (Hemiptera*: Cicadellidae) migration and severity of impact. PLoS One. 2015:10(5):e0124915. 10.1371/journal.pone.0124915.

Bandi V, Gutwin C. Interactive exploration of genomic conservation. In: Proceedings of Graphics Interface 2020 (GI’20). 2020.

Bartlett CR, Deitz LL, Dmitriev DA, Sanborn AF, Soulier-Perkins A, Wallace MS. The diversity of the true hoppers (Hemiptera: Auchenorrhyncha). In: Insect Biodiversity: Science and Society. Vol. II. 2018. p. 501–590. 10.1002/9781118945582.ch19.

Bruna T, Lomsadze A, Borodovsky M. GeneMark-EP+: eukaryotic gene prediction with self-training in the space of genes and proteins. NAR Genom Bioinform. 2020:2(2):lqaa026. 10.1093/nargab/lqaa026.

Bruna T, Lomsadze A, Borodovsky M. GeneMark-ETP significantly improves the accuracy of automatic annotation of large eukaryotic genomes. Genome Res. 2024:34(5):757–768. 10.1101/gr.278373.123.

Dudchenko O, Batra SS, Omer AD, Nyquist SK, Hoeger M, Durand NC, Shamim MS, Machol I, Lander ES, Aiden AP, et al. De novo assembly of the *Aedes aegypti genome usin*g Hi-C yields chromosome-length scaffolds. Science. 2017:356(6333):92–95. 10.1126/science.aal3327.

Dudchenko O, Shamim MS, Batra SS, Durand NC, Musial NT, Mostofa R, Pham M, Glenn St Hilaire B, Yao W, Stamenova E, et al. The Juicebox Assembly Tools module facilitates de novo assembly of mammalian genomes with chromosome-length scaffolds for under $1000. bioRxiv. 2018:254797. 10.1101/254797.

Durand NC, Robinson JT, Shamim MS, Machol I, Mesirov JP, Lander ES, Aiden EL. Juicebox provides a visualization system for Hi-C contact maps with unlimited zoom. Cell Syst. 2016:3(1):99–101. 10.1016/j.cels.2015.07.012.

Emms DM, Liu Y, Belcher L, Holmes J, Kelly S. OrthoFinder: scalable phylogenetic orthology inference for comparative genomics. bioRxiv. 2025:2025.07.15.664860. 10.1101/2025.07.15.664860.

Gabriel L, Bruna T, Hoff KJ, Ebel M, Lomsadze A, Borodovsky M, Stanke M. BRAKER3: fully automated genome annotation using RNA-Seq and protein evidence with GeneMark-ETP, AUGUSTUS and TSEBRA. Genome Res. 2024:34(5):769–777. 10.1101/gr.278090.123.

Gurevich A, Saveliev V, Vyahhi N, Tesler G. QUAST: quality assessment tool for genome assemblies. Bioinformatics. 2013:29(8):1072–1075. 10.1093/bioinformatics/btt086.

Hubley R, Smit A, Dfam Consortium. RepeatModeler2: de novo transposable element family identification and modeling. 2023 [accessed 2026 Jun 30]. https://github.com/Dfam-consortium/RepeatModeler.

Kabrick LR, Backus EA. Salivary deposits and plant damage associated with specific probing behaviors of the potato leafhopper, *Empoasca fabae, on alfalf*a stems. Entomol Exp Appl. 1990:56(3):287–304. 10.1111/j.1570-7458.1990.tb01407.x.

Koren S, Walenz BP, Berlin K, Miller JR, Bergman NH, Phillippy AM. Canu: scalable and accurate long-read assembly via adaptive k-mer weighting and repeat separation. Genome Res. 2017:27(5):722–736. 10.1101/gr.215087.116.

Lamp WO, Nielsen GR, Danielson SD. Patterns among host plants of potato leafhopper, Empoasca fabae (Homoptera: Cicadellidae). J Kans Entomol Soc. 1994:67(4):354–368. https://www.jstor.org/stable/25085541.

Li H, Handsaker B, Wysoker A, Fennell T, Ruan J, Homer N, Marth G, Abecasis G, Durbin R. The Sequence Alignment/Map format and SAMtools. Bioinformatics. 2009:25(16):2078–2079. 10.1093/bioinformatics/btp352.

Molligan J, Jacques J, Mukhopadhyay S, Pérez-López E. De novo assembly and annotation of the *Empoasca fabaemitochondri*al genome. Mitochondrial DNA B Resour. 2025:10(6):403–408. 10.1080/23802359.2025.2498740.

Plante N, Durivage J, Brochu AS, Dumonceaux T, Almeida Santos A, Torres D, Bahder B, Kits J, Dionne A, Légaré JP, et al. Leafhoppers as markers of the impact of climate change on agriculture. Cell Rep Sustain. 2024:1(2):100029. 10.1016/j.crsus.2024.100029.

Raupach K, Borgemeister C, Hommes M, Poehling HM, Sétamou M. Effect of temperature and host plants on the bionomics of *Empoasca decipiens (Homoptera:* Cicadellidae). Crop Prot. 2002:21(2):113–119. 10.1016/S0261-2194(01)00070-9.

Roach MJ, Schmidt SA, Borneman AR. Purge Haplotigs: allelic contig reassignment for third-gen diploid genome assemblies. BMC Bioinformatics. 2018:19(1):460. 10.1186/s12859-018-2485-7.

Ruan J, Li H. Fast and accurate long-read assembly with wtdbg2. Nat Methods. 2020:17(2):155–158. 10.1038/s41592-019-0669-3.

Santos AA, Vieira Araújo FH, Plante N, Siqueira da Silva R, Pérez-López E. Seasonal phenology of *Empoasca fabae(Hemiptera:* Cicadellidae) in Québec, Canada. Environ Entomol. 2025:54(5):1124–1135. 10.1093/ee/nvaf070.

Simão FA, Waterhouse RM, Ioannidis P, Kriventseva EV, Zdobnov EM. BUSCO: assessing genome assembly and annotation completeness with single-copy orthologs. Bioinformatics. 2015:31(19):3210–3212. 10.1093/bioinformatics/btv351.

Smit AFA, Hubley R, Green P. RepeatMasker Open-4.0. 2013 [accessed 2026 Jun 30]. http://www.repeatmasker.org.

Sun J, Lu F, Luo Y, Bie L, Xu L, Wang Y. OrthoVenn3: an integrated platform for exploring and visualizing orthologous data across genomes. Nucleic Acids Res. 2023:51(W1):W397–W403. 10.1093/nar/gkad313.

Taylor RAJ, Shields EJ. Revisiting potato leafhopper, *Empoasca fabae (Harris), m*igration: implications in a world where invasive insects are all too common. Am Entomol. 2018:64(1):44–51. 10.1093/ae/tmy009.

Vasimuddin M, Misra S, Li H, Aluru S. Efficient architecture-aware acceleration of BWA-MEM for multicore systems. In: 2019 IEEE International Parallel and Distributed Processing Symposium (IPDPS). 2019. p. 314–324. 10.1109/IPDPS.2019.00041.

Walker BJ, Abeel T, Shea T, Priest M, Abouelliel A, Sakthikumar S, Cuomo CA, Zeng Q, Wortman J, Young SK, et al. Pilon: an integrated tool for comprehensive microbial variant detection and genome assembly improvement. PLoS One. 2014:9(11):e112963. 10.1371/journal.pone.0112963.

Wang Y, Tang H, DeBarry JD, Tan X, Li J, Wang X, Lee T, Jin H, Marler B, Guo H, et al. MCScanX: a toolkit for detection and evolutionary analysis of gene synteny and collinearity. Nucleic Acids Res. 2012:40(7):e49. 10.1093/nar/gkr1293.

Zhang L, Wang F, Qiao L, Dietrich CH, Matsumura M, Qin D. Population structure and genetic differentiation of tea green leafhopper, *Empoasca (Matsumurasca) onukii, i*n China based on microsatellite markers. Sci Rep. 2019:9(1):1202. 10.1038/s41598-018-37881-0.

