## Supplementary material for "A Chromosome-level Genome Assembly of the Potato Leafhopper *Empoasca fabae* (Hemiptera: Cicadellidae)": Table S1 to S3

Supplementary Tables

Table S1. BUSCO completeness assessment of the *Empoasca fabae* reference genome.

| Metric | Value |
| --- | --- |
| Lineage dataset | hemiptera_odb10 |
| Total BUSCO groups searched (n) | 2,510 |
| Complete BUSCOs (C) | 2,317 (92.4%) |
| • Complete and single-copy (S) | 2,268 (90.4%) |
| • Complete and duplicated (D) | 49 (2.0%) |
| Fragmented BUSCOs (F) | 69 (2.7%) |
| Missing BUSCOs (M) | 124 (4.9%) |
| Internal stop codons detected | 76 of 2,317 (3.3%) |
| BUSCO version | 5.8.2 |

Table S2. Repeat composition of the *Empoasca fabae* genome.

| Category | Count | bp occupied | % of genome |
| --- | --- | --- | --- |
| Total sequences analyzed | 132 | — | — |
| Total genome length | — | 908,556,302 bp | — |
| Bases masked | — | 433,498,059 bp | 47.71% |
| Total interspersed repeats | — | 420,896,544 bp | 46.33% |
| Retroelements (total) | 416,321 | 140,008,245 bp | 15.41% |
| LINEs (total) | 154,445 | 57,140,554 bp | 6.29% |
| CRE/SLACS | 1,416 | 603,963 bp | 0.07% |
| L2/CR1/Rex | 99,983 | 35,840,522 bp | 3.94% |
| R1/LOA/Jockey | 2,362 | 1,267,584 bp | 0.14% |
| R2/R4/NeSL | 14,341 | 5,673,728 bp | 0.62% |
| RTE/Bov-B | 2,280 | 839,057 bp | 0.09% |
| L1/CIN4 | 0 | 0 bp | 0.00% |
| LTR elements (total) | 261,876 | 82,867,691 bp | 9.12% |
| BEL/Pao | 29,089 | 10,538,819 bp | 1.16% |
| Ty1/Copia | 17,525 | 5,567,510 bp | 0.61% |
| Gypsy/DIRS1 | 120,429 | 41,030,494 bp | 4.52% |
| Retroviral | 0 | 0 bp | 0.00% |
| DNA transposons (total) | 35,321 | 9,562,848 bp | 1.05% |
| hobo–Activator | 3,621 | 887,284 bp | 0.10% |
| Tc1–IS630–Pogo | 16,673 | 4,724,252 bp | 0.52% |
| En–Spm | 0 | 0 bp | 0.00% |
| MULE–MuDR | 111 | 53,352 bp | 0.01% |
| PiggyBac | 439 | 209,957 bp | 0.02% |
| Tourist/Harbinger | 8,713 | 1,353,394 bp | 0.15% |
| Other (Mirage, P-element, Transib) | 1,768 | 185,058 bp | 0.02% |
| Rolling-circle (Helitrons) | 283 | 144,488 bp | 0.02% |
| Unclassified repeats | 1,242,878 | 271,325,451 bp | 29.86% |
| Small RNAs | 30,072 | 6,879,898 bp | 0.76% |
| Satellites | 5 | 1,649 bp | 0.00% |
| Simple repeats | 103,811 | 4,626,206 bp | 0.51% |
| Low complexity regions | 19,917 | 949,274 bp | 0.10% |

Table S3. Female and male genomic coverage ratios across *Empoasca fabae* chromosomes.

| Chr | Length (bp) | Female Reads | Female per Mb | Male Reads | Male per Mb | F/M Ratio | Normalized (F/M ÷ 0.85) |
| --- | --- | --- | --- | --- | --- | --- | --- |
| 1 | 164,038,279 | 16,089,980 | 98,086 | 18,909,202 | 115,273 | 0.85 | 1.00 |
| 2 | 130,604,558 | 13,558,275 | 103,812 | 15,889,530 | 121,661 | 0.85 | 1.00 |
| 3 | 117,682,009 | 14,098,819 | 119,804 | 16,837,756 | 143,078 | 0.84 | 0.99 |
| 4 | 96,186,748 | 9,114,983 | 94,763 | 10,756,076 | 111,825 | 0.85 | 1.00 |
| 5 | 96,107,418 | 9,242,000 | 96,163 | 10,720,810 | 111,550 | 0.86 | 1.01 |
| 6 | 69,001,508 | 6,579,218 | 95,348 | 7,843,153 | 113,666 | 0.84 | 0.99 |
| 7 | 64,675,687 | 7,100,885 | 109,792 | 8,533,809 | 131,948 | 0.83 | 0.98 |
| 8 | 40,996,017 | 4,547,002 | 110,913 | 5,407,852 | 131,912 | 0.84 | 0.99 |
| 9 | 39,763,386 | 3,716,483 | 93,465 | 4,437,190 | 111,590 | 0.84 | 0.99 |
| X | 79,212,458 | 8,513,631 | 107,478 | 7,845,726 | 99,046 | 1.09 | 1.28 |
